# Effects of serotonergic psychedelics on synaptogenesis and immediate early genes expression – comparison with ketamine, fluoxetine and lithium

**DOI:** 10.1101/2024.08.07.606965

**Authors:** Yana Vella, Kateřina Syrová, Aneta Petrušková, Isis Koutrouli, Viera Kútna, Jan Pala, Marek Nikolič, Klára Šíchová, Vladimír Mazoch, Radek Jurok, Martin Kuchař, Zdeňka Bendová, Tomáš Páleníček

## Abstract

**Background:** Recent evidence suggests that psychedelics are able to induce rapid and long-lasting antidepressant effects. The generally acknowledged explanation for these traits is the phenomenon of neuroplasticity, although exact underlying molecular mechanisms remain unclear.

**Aims:** This study investigates selected neuroplastic effects of psilocin, lysergic acid diethylamide (LSD) and N,N-dimethyltryptamine (DMT) in direct comparison with ketamine, fluoxetine and lithium after acute (1 h) and/or prolonged (24 h) treatment *in vitro*.

**Methods:** Rat primary cortical cultures were treated with 10 µM psilocin, 1 µM lysergic acid diethylamide (LSD), 90 µM N, N-dimethyltryptamine (DMT), 1 µM ketamine, 10 µM fluoxetine and 5 mM lithium. Analysis of synaptic puncta was performed; puncta of presynaptic marker synapsin I/II, postsynaptic density protein 95 (PSD-95), and their co-localization (established synapse) were assessed 24 h after drug treatment. Next, expressions of immediate early genes (IEGs) encoding activity-regulated cytoskeleton-associated protein (*Arc*), early growth response 1 (*Egr1*), and neuronal PAS (Per-ArntSim) domain protein 4 (*Npas4*) were analysed 1 and 24 h after drug treatments.

**Results:** Psilocin increased synaptic puncta count and induced *Arc* expression. The effect to promote synaptogenesis was comparable to ketamine and lithium; ketamine additionally increased PSD-95 puncta count. LSD and DMT didn’t induce any significant effect. Interestingly, fluoxetine had no effect on synaptic puncta count, but upregulated *Egr1* and *Npas4*.

**Conclusions:** Psilocin demonstrated a significant neuroplastic effect comparable to that of ketamine and lithium, adding another piece of evidence to its profile as a promising therapeutic agent.

## Introduction

After years of complete marginalization of classic psychedelics, research on their pharmacology is gaining robust support regarding their therapeutic potential on a plethora of neuropsychiatric conditions including depression, anxiety, post-traumatic stress disorder, addiction, anorexia, or chronic pain (Cameron et al., 2023; Carhart-Harris & Goodwin, 2017). As a positive control, psychedelic studies often comprise ketamine, a dissociative anaesthetic which produces psychoactive effects in subanaesthetic doses and has a confirmed efficacy as a fast-acting antidepressant. While classic psychedelics and ketamine aim at distinct receptor systems (serotonergic and glutamatergic, respectively) they share significant features beyond being psychoactive; they rapidly increase dendritogenesis, spinogenesis and synaptogenesis (De la Fuente Revenga et al., 2021; Li et al., 2010; Ly et al., 2021; Moliner et al., 2023; Raval et al., 2021), and induce a downstream network reconfiguration through complex neuroplastic mechanisms, hence the term “psychoplastogens” has recently been proposed (Ly et al., 2018).

A growing body of research postulates a correlation between many neuropsychiatric disorders and disrupted neuroplasticity, accompanied by an atrophy of neurons in the cortical and limbic brain regions (Rădulescu et al., 2021; Soeiro-de-Souza et al., 2012). Vice versa, findings have demonstrated that antidepressant drugs, including selective reuptake inhibitors (SSRIs), exert their therapeutic effects by reactivating a plasticity window in the adult cortex, which resembles juvenile-like brain processes. In such cases, neuronal networks are being shaped by sensory information through experience, by regulating the expression of genes related to plasticity and resilience, among other mechanisms (Puglisi-Allegra et al., 2021; Umemori et al., 2018). Although these medicaments have been the mainstay of treatment for depression, they possess several drawbacks; the antidepressant effects take 6-8 weeks of adequate doses to become evident, various side effects can appear, and many patients are unresponsive to this first line treatment. In line with the delayed onset of antidepressant actions, the neuroplastic changes take place only after chronic administration.

Lithium is a drug primarily used as a mood stabilizer in bipolar disorder, but also as an augmentation strategy option in the treatment of depression. Recent evidence confirms its robust neuroplastic effects, such as neuroprotection, glioprotection, elevated adult neurogenesis and synaptic plasticity (Hammonds & Shim, 2009). It has been shown to elevate postsynaptic density protein 95 (PSD-95) in rat cortical and hippocampal cultures (Kim & Thayer, 2009; Tai et al., 2022). In spite of that, use of lithium has some limitations considering its possible adverse effects (Antoniadou et al., 2018; Cameron et al., 2023).

In contrast to common antidepressants, psychedelics and ketamine present a robust and rapid therapeutic efficacy even in treatment-resistant depression patients, most likely by the increase of structural and functional plasticity in the prefrontal cortex (PFC), hippocampus, and other brain regions involved in emotions (Krystal et al., 2019; Vargas et al., 2023). Depending on the drug and dosage regimen, sustained antidepressant effects are observed.

Immediate early genes (IEGS) are a family of genes that are rapidly and transiently transcribed after a wide variety of signals. As opposed to the “late response” genes, IEGs expression does not depend on any precedent protein synthesis. They encode many functionally distinct proteins, such as transcription and growth factors, cytoskeletal proteins or signalling molecules, and play an essential role in cellular responses that contribute to long-term neuronal plasticity.

*Arc*, *Egr1* and *Npas4* are three IEGs linked to synaptic plasticity processes. Arc is a postsynaptic effector protein localised primarily in the dendrites of excitatory glutamatergic neurons of cortex and hippocampus. Arc alters spine morphology and synaptic strength via α-amino-3-hydroxy-5-methyl-4-isoxazolepropionic acid receptor (AMPAR) internalisation, and therefore has a key role in synaptic plasticity and long-term memory. Psychedelics have been found to induce its expression (Nichols & Sanders-Bush, 2002; Pei et al., 2000). Egr1 is a transcription factor which, following synaptic activation, binds to the *Arc* promoter and activates its transcription in a non-dependent manner. Egr1 is a transcription suppressor of PSD-95 and modulates N-methyl-D-aspartate receptor (NMDAR)-dependent AMPAR trafficking (Qin et al., 2015). Egr1 is specifically involved in memory encoding and synaptic organization at multiple levels, resulting in synaptic stabilization. Increase in *Egr1* gene expression in murine cortex has been found after LSD administration, as well as after fluoxetine *in vitro* (González-Maeso et al., 2003, 2007; Levy et al., 2019). Npas4 is a transcription factor selectively activated by neuronal activity, directly regulating brain-derived neurotrophic factor expression (Sun & Lin, 2016).

The exact pathways activated by psychedelics have been under intense investigation in recent years. Despite that, to the best of our knowledge, no study has directly compared these promising agents to regular psychiatric medication in terms of their neuroplastic effects. In this study, psilocin, DMT and LSD were examined and compared to two of the most prescribed psychiatric drugs, fluoxetine and lithium, as well as with ketamine, a generally acknowledged psychoplastogen, in the ability to promote synaptogenesis and to enhance the gene expression of selected IEGs *Arc*, *Egr1* and *Npas4*.

## Methods

### Primary cortical culture isolation and maintenance

All experiments were conducted in accordance with guidelines of the Directive 2010/63/EU of the European Parliament and approved by the Animal Care and Use Committee of the Ministry of Health (reference number MZDR 17637/2020-4/OVZ) in line with the Animal Protection Code of the Czech Republic 246/1992. NIMH is authorized for the use of experimental animals (accreditation ref. no. MZE-51088/2021-13114).

Primary cortical cultures were prepared as previously described by Jorratt et al. (2023) with minor modifications. Briefly, embryos were extracted from time-mated female Wistar rats on embryonic day 18. The cortices were dissociated under the stereomicroscope VisiScope® 250 (VWR, USA) and put in ice-cold Dulbecco’s phosphate-buffered saline (DPBS, Gibco, 14190144). Then, the cortices were washed thrice with ice-cold DPBS and mechanically dissociated with a 0.9 mm needle (B. Braun, 4657519), followed by a 0.45 mm needle (B. Braun, 4657683), each needle thrice. The dissociated supernatant was transferred into a seeding medium containing advanced Dulbecco’s modified Earle’s medium (DMEM, Gibco, 12491015) supplemented with 10% fetal bovine serum (FBS, Biowest, S1810-500), 1% Antibiotic-Antimycotic solution (Gibco, 15240062) and 1% GlutaMAX supplement (Gibco, 35050061), and filtered twice through a 100-µm cell strainer (VWR, 732–2759). The singularized cells were counted using a haemocytometer and seeded on poly-L-lysine (Merck, P1274-25MG) pre-coated well plates. 24-well plates with inserted 13-mm, #1.5 coverslips (VWR, 631-0150) with a seeding density of 50,000 cells/well were used for immunocytochemistry (ICC), 6-well plates with a density of 500,000 cells/well were used for real-time quantitative polymerase chain reaction (qPCR). The cells were cultured in a humidified incubator at 37°C and 5% CO_2_. On the next day, the seeding medium was replaced with neurobasal medium (Gibco, 21103049), enriched with 2% B27 supplement (Gibco, 17504044), 1% Antibiotic-antimycotic and 1% GlutaMAX. Half of the growth medium was changed twice a week.

### Drugs and chemicals

On day *in vitro* (DIV) 17, the cultures were treated for 24 h for the ICC analysis of colocalization, and for 1 h and 24 h for the qPCR analysis of changes in IEGs expression. The cells were treated with 10 µM psilocin (THC Pharm, Germany), 1 µM LSD fumarate (Alfarma, Czechia), 90 µM DMT fumarate (Forensic Laboratory of Biologically Active Substances, University of Chemistry and Technology, Czech Republic), 10 µM fluoxetine hydrochloride (Molekula Group, Czechia), 5 mM lithium chloride (Merck, Germany) and 1 µM ketamine (Merck, Germany). All drugs were of ≥ 99% purity and were diluted in sterile distilled water. Sterile distilled water was used as negative control (Ctrl).

### Immunocytochemistry

Cells were quickly washed with pre-warmed (37°C) phosphate buffer saline (PBS) and fixed with pre-warmed 4% paraformaldehyde in PBS for 4 min, then rinsed 3 x 5 min with PBS. Next, blocking and permeabilization were done in one step with 10% FBS, 0.1% Triton X-100 in PBS for 45 min at RT. The coverslips were subsequently incubated with primary antibodies chicken anti-MAP2 (1:10,000, Abcam, ab5392), mouse anti-PSD-95 (1:500, Abcam, ab192757), and guinea-pig anti-synapsin I/II (1:1000, Synaptic systems, 106004) at 4°C overnight. The following day, coverslips were washed 3 x 5 min with PBS and incubated with secondary antibodies AlexaFluor (AF) 647 conjugated donkey anti-chicken (703-605-155), AF488 donkey anti-guinea pig (706-545148) and AF594 donkey anti-mouse (715-545-151) for 1 h at room temperature in the dark. All secondary antibodies were purchased from Jackson ImmunoResearch and diluted 1:500 for working solutions. After the incubation, coverslips were washed 3 x 5 min with PBS in the dark and rinsed with distilled water, dried from excess liquid, mounted onto microscope slides using ProLong™ Gold Antifade Mountant with DAPI (Invitrogen, P36941), and allowed to cure overnight. All the antibodies were diluted in the staining solution consisting of 0.1% Triton X-100 in PBS.

### Image acquisition and analysis

Images were acquired with confocal laser scanning microscope Leica SP8 X on inverted microscope scope Leica DMi8 (Leica Microsystems, Germany). Plan apochromat oil objective Leica 63x/1.40 CS2 was used. The confocal pinhole was set to Airy1, and bidirectional scanning with line averaging 4 times was applied for Z-stacks at 0.3 μm Z-step size. The acquisition was done in 8-bit, 1024×1024 pixels, and 2.6x digital zoom. For every experiment, samples were imaged under identical software settings. Automatic threshold selection was performed, and binary pictures were analysed using the Synapse Counter plugin (Dzyubenko et al., 2016) for Fiji (version 1.52e, Schindelin et al., 2012), following a protocol from Li et al. (2016).

### Changes in IEGs expression

Cells were lysed in RNA Blue reagent (TopBio, R013) and phenol-chloroform-based extraction of the homogenate was performed according to the manufacturer’s instructions. Complementary DNA (cDNA) was generated using the same amount of each total RNA sample within one batch. High-capacity cDNA Reverse Transcription Kit (Applied Biosystems, 4368814) was used according to the manufacturer’s instructions. qPCR was performed with CFX96 Touch Real-Time PCR Detection System (Bio-Rad Laboratories, USA) using Luna Universal qPCR Master Mix (New England Biolabs, M3003L) according to the manufacturer’s instructions. IEGs signal was normalized to housekeeping genes glyceraldehyde-3-phosphate dehydrogenase (*Gapdh*) and peptidylprolyl isomerase A (*Ppia*), which were selected as an optimal reference via NormFinder tool (version 0.953, Andersen et al., 2004). Comparative Ct (ΔΔCt) method was used to determine the changes in mRNA levels of *Arc*, *Egr1*, and *Npas4* (Livak & Schmittgen, 2001).

The primers were designed in the NCBI Primer-BLAST tool (Ye et al., 2012) or selected based on literature (Adams et al., 2017; Heroux et al., 2018) and the sequences were chosen as follows: *Arc* (5’- GCAGAAAGCGCTTGAACTTG-3’, 5’-AGCGGGACCTGTACCAGAC-3’), *Egr1* (5’- CATGCAGATTCGACACTGGAAG-3’, 5’-GTATGCTTGCCCTGTTGAGTCC-3’), *Npas4* (5’- GCCACTATGTCTTCAAGCTCT-3’, 5’-CTGCATCTACACTCGCAAGG-3’), *Gapdh* (5’- GGCATGGACTGTGGTCATGA-3’, 5’-CAACTCCCTCAAGATTGTCAGC-3’), *Ppia* (5’- TCCTTTCTCCCCAGTGCTCAG-3’, 5’-TCAACCCCACCGTGTTCTTC-3’).

### Data and statistical analysis

Statistical analyses were performed in GraphPad Prism (version 8.0.1, GraphPad Software, Inc., USA). Outliers were removed using the ROUT method with a 1 % Q value. Data violating the assumption of normality (Shapiro-Wilk test) were normalised by square root transformation. Homogeneity of variance was tested using the Brown-Forsythe test. One-way analysis of variance (ANOVA) with Dunnett’s *post hoc* test were performed for the comparison of drug treatment effect compared to NC.

## Results

### Psilocin induces synaptogenesis

Synaptic puncta were identified by the co-localization of synapsin I/II and PSD-95 puncta. Analysis of changes in synapsin I/II density did not show any effect of the treatment (Fig. 1A). Significant changes were observed in PSD-95 puncta density after the treatment [F (6, 330) = 5.55, p < 0.001], *post hoc* analysis revealed prominent effect of ketamine (p < 0.001, Fig. 1B). Analysis of co-localized (i. e. synaptic) puncta also displayed significant effect of the drug treatment [F (6, 333) = 8.60, p < 0.001], further *post hoc* revealed strong effect of psilocin, lithium, and ketamine (all p < 0.001, Fig. 1C).

**Figure 1.**
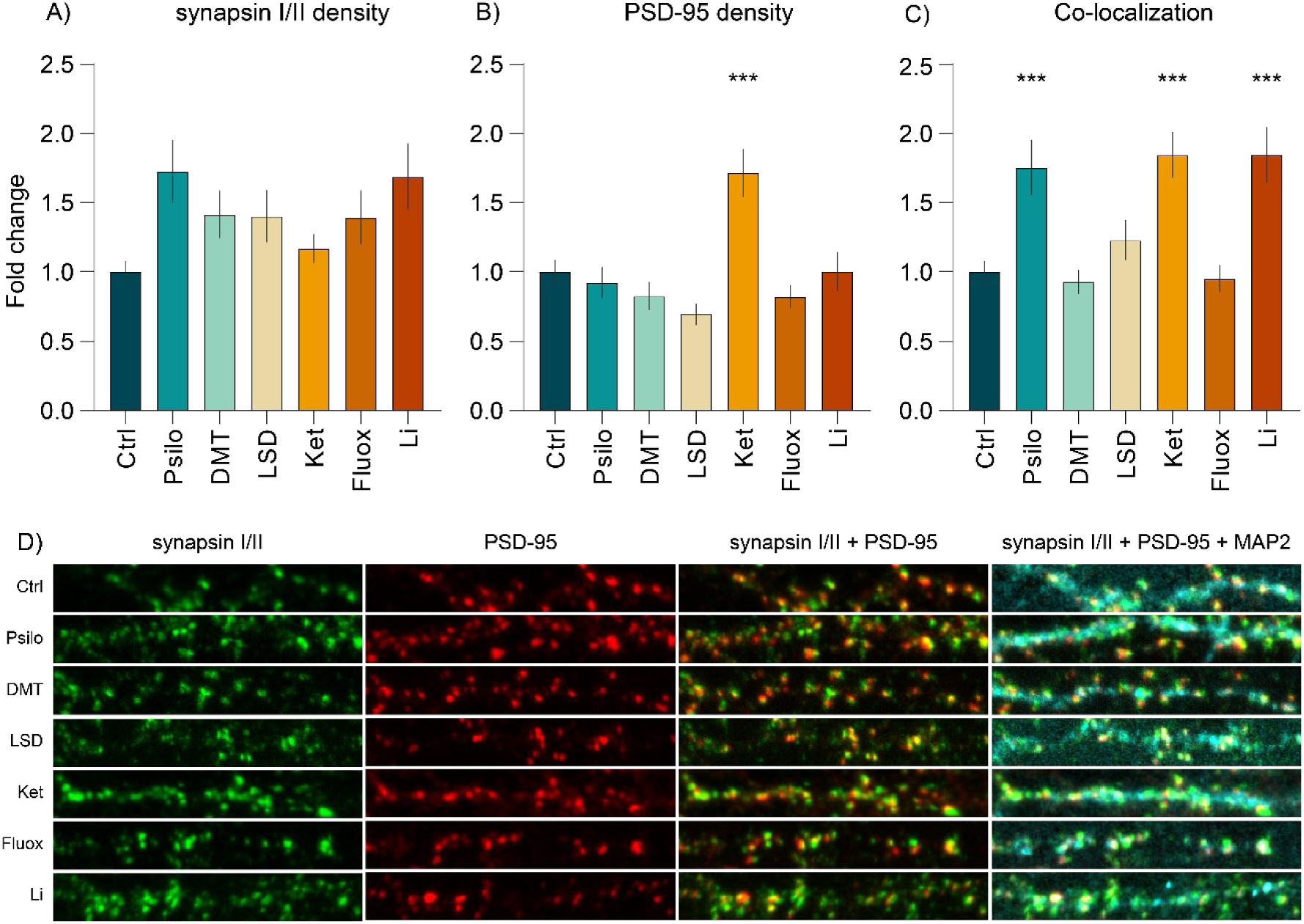
The effect of selected psychoplastogens on synaptogenesis. Quantification of presynaptic density (**A**, synapsin puncta), postsynaptic density (**B**, PSD-95 puncta) and their co-localization (**C**, established synapses) after 24 h drug treatment of primary cortical neurons. n = 38–79 randomly chosen neurons per treatment from at least three biological replicates. One-way ANOVA followed by Dunnett’s *post hoc* test was performed, data are presented as mean fold change ± SEM. *p < 0.05, **p < 0.01, ***p < 0.001. **D** Representative images of primary dendrites of cortical neurons (DIV18) after 24 h treatment. Orange areas in the synapsin + PSD-95 merged images indicate colocalization. Green, synapsin I/II; red, PSD-95; cyan, MAP2.

### Psilocin induces gene expression of *Arc* after acute treatment

After one hour (Fig. 2A), *Arc* expression was significantly affected by the treatment [F (6, 24) = 4.73, p < 0.01], *post-hoc* analysis showed significant effect of psilocin (p < 0.01). Expression of *Npas4* was also significantly changed after one hour [F (6, 26) = 2.55, p < 0.05], although *post hoc* test did not reveal any specific drug treatment effect.

**Figure 2.**
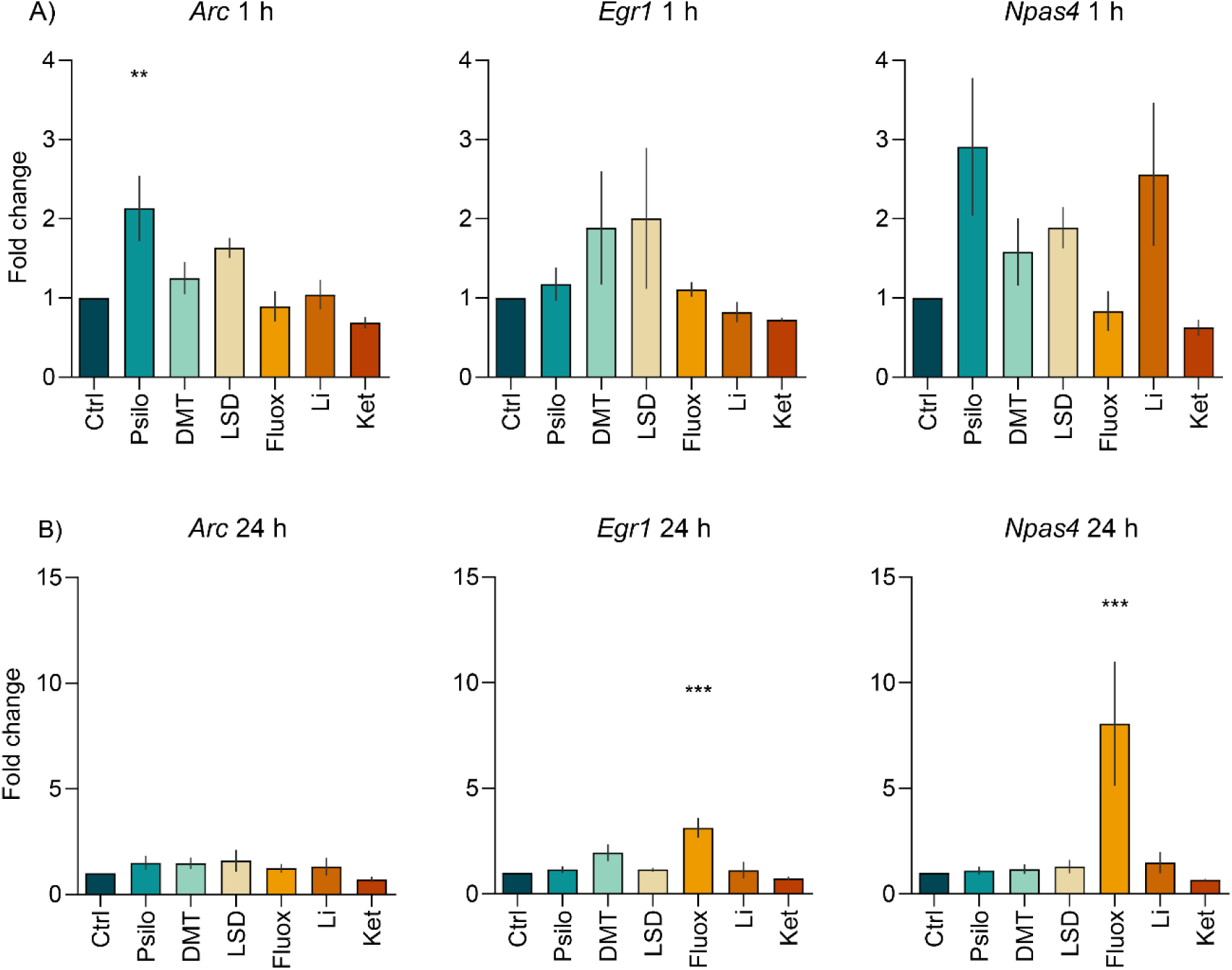
Effect of selected psychoplastogens on the expression of IEG mRNA. **A**+**B** Assessment of the changes in mRNA expression of IEG *Arc*, *Egr1* and *Npas4* after 1 h (**A**) and 24 h (**B**) drug treatment of primary cortical neurons. Data were collected from at least four biological replicates. One-way ANOVA followed by Dunnett’s *post hoc* test was performed. Data are presented as mean fold change ± SEM. *p < 0.05, **p < 0.01, ***p < 0.001.

After 24 hours (Fig. 2B), *Egr1* levels were significantly affected by the treatment [F (6, 21) = 8.34, p < 0.001], *post hoc* test showed significant effect of fluoxetine (p < 0.001). *Npas4* expression was also significantly altered by the treatment [F (6, 20) = 7.59, p < 0.001], *post hoc* test revealed prominent changes following fluoxetine treatment (p < 0.001).

## Discussion

The main finding of our study is that psilocin significantly promotes synaptogenesis and expression of the IEG *Arc*, and moreover, the effects of psilocin were comparable to those of lithium and ketamine. Surprisingly, we did not observe such effects after LSD and DMT treatment. Concerning the positive control treatments, ketamine also increased PSD-95 density and fluoxetine did not affect synaptic density, but upregulated *Egr1* and *Npas4* expression.

Our finding that psilocin increases neuronal plasticity is in line with other studies (Du et al., 2023; Moliner et al., 2023; Raval et al., 2021; Vargas et al., 2023). In accordance with Ly et al. (2018), we did not observe any effect on PSD-95 density after the treatment with DMT, but we discovered the increased PSD-95 and synapse densities associated with ketamine treatment, which was described also in other studies (Li et al., 2010; Ly et al., 2021). Interestingly, (Ly et al., 2018, 2021) observed also the effect of LSD on presynaptic and synaptic puncta count, as opposed to our results. The fact that we did not find any significant effect of LSD and DMT on synaptogenesis was unexpected. The explanation could be that we did not add dimethyl sulfoxide (DMSO) into the drug solution for the elimination of any noxious effect on the primary cells. However, since DMSO is a permeation enhancer, its lack in the solution might cause the non-significant results for LSD and DMT in our study.

In the study of changes in IEGs mRNA expression after psychedelic treatment, we found a significant upregulation of *Arc* after the acute treatment with psilocin, and of *Egr1* and *Npas4* following the prolonged treatment with fluoxetine. In contrast to our study, 90 min psilocybin treatment did not exhibit any significant changes in *Arc* in the PFC, but decreased expression in the hippocampi of rats (Jefsen et al., 2021). Another animal study demonstrates upregulated *Arc* expression following 90 min LSD treatment in PFC, but not in hippocampus (Nichols & Sanders-Bush, 2002). Davoudian et al. (2023) describe comparable elevation of IEG *cFos* in many brain regions after psilocybin and ketamine administration, with selected brain regions manifesting drug-preferential differences. Concerning *Egr1*, it has been shown that an acute dose of psychedelics increases its expression (Desouza et al., 2021; Liu et al., 2023), while chronic microdosing does not (Cameron et al., 2019). A study using noribogaine observed upregulated *Egr1* and *Npas4* expression in male mice with serotonin 2A receptor (5-HT2AR) knockout, *Npas4* expression in 5-HT2AR knockout females, and *Egr1* expression in wild type females; indicating sex and genotype differences in IEGs expression (Villalba et al., 2024). In this study, *Npas4* was clearly upregulated in a 5-HT2AR activation-independent manner. On top of that, noribogaine has a polypharmacological profile and does not aim directly at 5-HT2AR, as opposed to psychedelics. Another reason for the divergent results with aforementioned studies could dwell in our *in vitro* model, which comprises the mixture of cortical cells from both male and female embryos. This aspect may be necessary to take into account for the following studies focusing on IEGs, as it could be the cause of major variances.

Overall, our study reveals that psilocin indeed acts as a neuroplastic agent by enhancing both synaptogenesis and *Arc* expression. The co-localization results also indicate similar mechanisms by which psilocin, ketamine and lithium promote their neuroplastic effect on the synapse. On the other hand, our findings also suggest that the effects of fluoxetine on neuroplasticity are produced through different mechanisms than those of serotonergic psychedelics or ketamine.

## Acknowledgements

This work was supported by a grant from the Czech Science Foundation (project 20-25349S), Czech Health Research Council (project NU21-04-00307), Long-term conceptual development of research organization (RVO 00023752), Ministry of the interior of the Czech Republic (project VK01010212), Specific University Research, Czech Ministry of Education, Youth and Sports (project 260648/SVV/2024), ERDF-Project Brain dynamics, No. CZ.02.01.01/00/22_008/0004643, Charles University research program Cooperatio-Neurosciences, Grant Agency of Charles University in Prague, Czech Republic (project GAUK 430122) and private funds obtained via PSYRES - Psychedelic Research Foundation (psyresfoundation.eu).

## Author contributions

All authors made a significant contribution to the design, acquisition, analysis, or interpretation of data for this study. AP and TP designed the project, YV and KS carried out the experiments, YV, KS and IK wrote the manuscript. VK performed the dissections and supervised the ICC assay, KŠ, VM and MN carried out the statistical analyses, JP trained the experimenters in confocal microscopy and supervised the setup, MK and RJ synthesized and provided the group with ketamine, TP and ZB supervised the project and manuscript writing. All authors gave final approval for the current version of this work to be published. All authors agree to be accountable for all aspects of the work in ensuring that questions related to the accuracy or integrity of any part of the work are appropriately investigated and resolved.

## Data Availability

All data supporting described findings can be obtained from the corresponding author upon reasonable request.

